# High-throughput fluorescence-based screen identifies the neuronal microRNA miR-124 as a positive regulator of alphavirus infection

**DOI:** 10.1101/758201

**Authors:** Paula López, Erika Girardi, Bryan C. Mounce, Amélie Weiss, Béatrice Chane-Woon-Ming, Mélanie Messmer, Pasi Kaukinen, Arnaud Kopp, Diane Bortolamiol-Becet, Ali Fendri, Marco Vignuzzi, Laurent Brino, Sébastien Pfeffer

**Affiliations:** Architecture et Réactivité de l’ARN, Université de Strasbourg, Institut de Biologie Moléculaire et Cellulaire du CNRS, 67084 Strasbourg, France; Viral Populations and Pathogenesis Unit, Department of Virology, Institut Pasteur, CNRS UMR 3569, Paris, France; Institut de Génétique et Biologie Moléculaire et Cellulaire, 67400 Illkirch-Graffenstaden, France; Department of Microbiology and Immunology, Stritch School of Medicine, Loyola University Chicago, Maywood, IL 60153, USA

**Keywords:** microRNA, miR-124, RNA virus, arbovirus, alphavirus, Sindbis virus, chikungunya virus, neuron, host-virus interaction

## Abstract

Micro (mi)RNAs are small regulatory RNAs, which act as guide for the RISC complex to modulate the expression of target genes. In addition to their role in maintaining essential physiological functions in the cell, miRNAs can also regulate viral infections. They can do so directly by targeting RNAs of viral origin or indirectly by targeting RNAs from the host and this can result in a positive or negative outcome for the virus. Here, we performed a miRNA genome-wide screen in order to identify cellular miRNAs involved in the regulation of arbovirus infection in human cells. We identified sixteen miRNAs showing a positive effect on the virus, among which a number of neuron-specific ones such as miR-124. We confirmed that overexpression of miR-124 increases Sindbis virus (SINV) viral production and that this effect is mediated by its seed sequence. We further demonstrated that the SINV genome possesses a binding site for miR-124-3p. Both inhibition of miR-124-3p or silent mutations to disrupt this binding site in the viral RNA abolished the positive regulation. We also proved that miR-124 inhibition reduces SINV infection in human differentiated neuronal cells. Finally, we showed that the proviral effect of miR-124 is conserved for other medically relevant alphaviruses. Indeed, inhibition of miR-124 expression reduces chikungunya virus (CHIKV) viral production in human cells. Altogether, our work expands the panel of positive regulation of the viral cycle by direct binding of host miRNAs to the viral RNA and provides new insights into the role of cellular miRNAs as regulators of alphavirus infection.

**SIGNIFICANCE STATEMENT:** Arthropod-borne (arbo) viruses are part of a class of pathogens that are transmitted to their final hosts by insects. Because of climate change, the habitat of some of these insects, such as mosquitoes, is shifting, thereby facilitating the emergence of viral epidemics. Among the pathologies associated with arboviruses infection, neurological diseases like meningitis or encephalitis represent a significant health burden. Using a genome-wide miRNA screen, we identified the neuronal miR-124 as a positive regulator of the Sindbis and chikungunya alphaviruses. We also showed that this effect was in part direct, thereby opening novel avenues to treat alphaviruses infection.

## INTRODUCTION

Infectious diseases, and among them viral diseases, remain a leading cause of morbidity and mortality worldwide. In addition to the direct consequences of viral infections, many health disorders are indirectly linked to viruses. Although vaccines are available for some viruses, this is not the case for a large number of them, and it is essential to find novel antiviral compounds to fight them. Alphaviruses, from the *Togaviridae* family, are arthropod-borne viruses (arboviruses) transmitted to vertebrates by a mosquito vector and form a group of widely distributed human and animal pathogens. They are small, enveloped, positive single-stranded RNA viruses. Their RNA genome of ∼11kb is capped and polyadenylated. It has two open reading frames (ORFs) encoding nonstructural and structural proteins. ORF2 is expressed through the production of a sub-genomic RNA from an internal promoter in the minus-strand RNA replication intermediate. In addition to the protein-coding sequences, alphavirus RNAs contain important regulatory structures, such as the 5′ and 3′ UTRs (1).

Alphaviruses represent an emerging public health threat as they can induce febrile and arthritogenic diseases, as well as other highly debilitating diseases, such as encephalitis (2). Sindbis virus (SINV) is considered as the prototypical alphavirus and is widely used as a laboratory model. Although the infection has been mainly associated to rash, arthritis and myalgia in humans (3), SINV displays a neuronal tropism in developing rodent brain cells and is associated with encephalomyelitis (4). Another virus from the same genus is chikungunya virus (CHIKV), which causes outbreaks of severe acute and chronic rheumatic diseases in humans (5). CHIKV has also been reported to affect the human nervous system causing encephalopathy in newborns, infants and adults (6). Due to its ability to quickly spread into new regions, CHIKV is classified as an emergent virus (7, 8), for which preventive or curative antiviral strategies are needed.

Micro (mi)RNAs are small 22 nucleotide long non-coding RNAs, which act as guides for effector proteins to post-transcriptionally regulate the expression of target cellular mRNAs (9) but also viral RNAs (10–17). These small RNAs have been identified in almost all eukaryotic species, and a number of them are conserved throughout evolution (18). They derive from longer precursors, which are transcribed by RNA polymerase II, and are sequentially processed by the RNase III enzymes Drosha and Dicer. The mature miRNA is then assembled in a protein of the Argonaute family, to guide it to target RNAs. Once bound to its target, the Argonaute protein regulates its expression by recruiting proteins to inhibit translation initiation and induce its destabilization by deadenylation (19). The main determinant of miRNA sequence specificity is its seed sequence which corresponds to a short region at the 5′-end of miRNAs (nucleotides 2–7) (20). Perfect pairing of the miRNA seed with the target RNA represents the minimal requirement for efficient Argonaute binding and function. In some cases, additional base pairing toward the 3′ end of the mature miRNA (so-called 3′-compensatory sites) may compensate for suboptimal pairing in the seed region (21, 22).

Identifying miRNAs that alter virus replication has illuminated roles for these molecules in virus replication and highlighted therapeutic opportunities. Target predictions based on the concept of “seed” initially identified binding sites for the liver-specific miR-122 in the 5’UTR of hepatitis C virus (HCV), which turned out to be positively regulated by this miRNA (23). Further work from different teams later showed that miR-122 can positively regulate the virus by increasing stability and translation of the viral RNA (24, 25). Interestingly, the use of inhibitors of miR-122 in HCV infection is currently in clinical trial as a miRNA-based treatment for antiviral therapy (26).

Here, we performed a genome-wide miRNA overexpression and inhibition screen to identify cellular miRNAs involved in the regulation of SINV infection. We found 16 miRNAs with a positive effect on virus accumulation. Among them, we showed that miR-124-3p positively regulates the virus, and we identified a binding site for this small RNA in the viral genome. miR-124 is the most expressed, conserved and specific microRNA in the central nervous system (CNS) and it has been described as a master regulator of neuronal differentiation (27). It is up-regulated as neuronal progenitors exit mitosis and begin to differentiate (28) and its expression has been shown to be sufficient to drive cells into the neuronal pathway (29). Mutations in the miR-124-3p seed region or in the binding site on the viral RNA abolished the regulation. We also proved that the miR-124-3p inhibition strategy *via* antisense oligonucleotides reduces both SINV and CHIKV production in human cells expressing this miRNA. These studies highlight a novel role for miR-124-3p in arbovirus infection and suggest targeting alphavirus infection *via* miRNA modulation could be used therapeutically.

## RESULTS

### Identification of miRNAs involved in the regulation of SINV infection

To identify miRNAs that are involved in the regulation of SINV infection in a systematic and comprehensive way, we performed a fluorescence microscopy-based, high-throughput screening using two libraries of respectively ∼2000 human miRNA mimics and ∼2000 antimiRNA oligonucleotides (Figure 1A). Control miRNA mimics or antimiRNAs with no sequence homology to any known human miRNA were used as negative controls. A short interfering RNA (siRNA) against the 3’ UTR of SINV was used as a positive control. Huh7.5.1 cells were transfected with either the library of miRNA mimics or antimiRNAs and 72 hours later the cells were infected with a SINV strain encoding the green fluorescent protein (GFP) (Figure S1). The number of GFP positive cells and the GFP intensity were measured 24 hours post-infection (hpi) by automated image analysis and the robust Strictly Standardized Mean Difference (SSMD* or SSMDr) value was calculated for each miRNA to identify significant hits (Figure 1B-D). While the antimiRNA screen did not reveal any significant hit, the mimic screen identified 16 miRNAs that increased the GFP signal significantly (SSMDr >1.28) without affecting cell viability (Figure 1B-C). All of the top hit miRNAs were undetectable or expressed at a very low level in Huh7.5.1 cells as assessed by small RNA sequencing (Figure 1C, right column and Figure S2A), which explains why transfection of the corresponding antimiRNAs showed no effect on GFP levels. Among the top hits, miR-124-3p and miR-129-5p which are known to be expressed in neuronal cells (30, 31) (Figure S2B-C), had the most striking effect (SSMDr > 2.20). Moreover, we could show that other hits such as miR-1244, miR-665 and miR-30b-3p were enriched in mouse neuronal tissues compared to other tissues such as heart or liver (Figure S2C), suggesting a link between neuron-enriched miRNAs and positive regulation of the virus.

**Figure 1.**
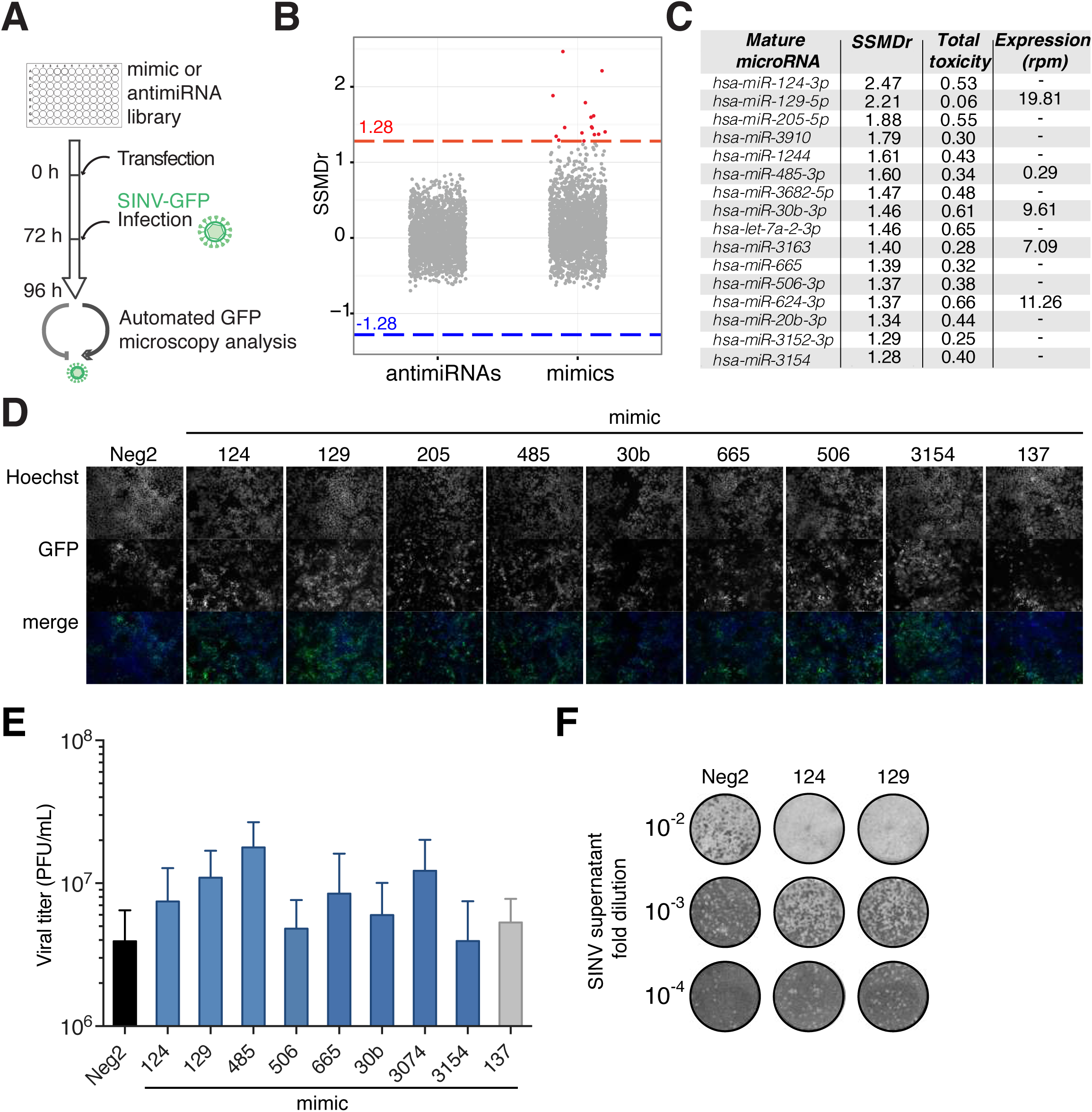
Identification of cellular miRNAs positively regulating SINV infection in Huh7.5.1 cells. (**A**) Schematic representation of the screening protocol. Huh7.5.1 cells were reverse transfected with a complete library of mimics or antimiRNAs corresponding to all identified human miRNAs and 72 h post-transfection they were infected with SINV-GFP at a low multiplicity of infection (MOI 10^−3^) for 24 h. The infection level was analyzed by automated microscopy analysis based on GFP fluorescence. (**B**) Dot plot representation of the strictly standardized mean difference (SSMDr) in fluorescence for each mimic or antimiRNA. Candidates with an absolute SSMDr equal or higher than the threshold 1.28 were considered as significant. In red, significant mimic candidates. (**C**) List of significant candidates identified in the mimic screen with their respective SSMDr and associated toxicity values. miRNA expression levels in Huh7.5.1 cells determined by small RNA-seq are indicated as read per million miRNA reads (rpm). (**D**) Representative fluorescence images from the mimic screen of some of the top candidates compared to the controls (Neg#2 and miR-137). The top panel corresponds to Hoechst staining of cell nucleus, the middle panel corresponds to GFP signal from infected cells and the lower panel is the merge. (**E**) Viral titers produced by SINV-GFP after transfection of top candidates (blue bars) in Huh7.5.1 cells compared to control mimic (black bar) and negative control miR-137 (grey). PFU: plaque forming units. Error bars represent mean ± SEM of three independent experiments. (**F**) Representative plaque assay image of SINV infected (at indicated MOIs) Huh7.5.1 cells transfected with control or candidate mimics.

To determine whether the positive effect of these miRNAs on SINV-GFP expression implied an increase in viral production, we measured the effect of the overexpression of selected miRNAs on SINV titers in two different cell types compared to a negative miRNA mimic control (cel-miR-67) or to a miRNA whose expression did not alter GFP levels in the screen (hsa-miR-137) (Figure 1E and S3). Similarly to the GFP fluorescence, all of the eight miRNAs tested (namely miR-124-3p, miR-129-5p, miR-485-3p, miR-506-3p, miR-665, miR-30b-3p, miR-3074, and miR-3154) increased SINV-GFP viral titers in Huh7.5.1 cells, providing strong evidence for their involvement in the positive regulation of viral infection (Figure 1E and F). Moreover, overexpression of miR-124-3p, miR-129-5p, miR-205-5p, miR-485-3p, miR-506-3p and miR-3154 showed a proviral effect on the virus in HEK293A cells as well (Figure S3). Interestingly, among the top candidates we found miR-124-3p and miR-506-3p showing a proviral effect in the screen. These miRNAs both belong to the same microRNA family and share similar physiological functions (32), which consolidates our method and suggests strong evidence for a possible implication of the seed region in the regulation of SINV infection.

### Mutations in miR-124-3p seed or in its binding site on the viral RNA abolish the positive regulation on SINV

Because miR-124 is the most expressed miRNA in neuronal tissues, we decided to focus our efforts on the characterization of the mechanism underlying its effect on SINV. In order to assess whether the proviral effect observed depended on the miRNA seed sequence, we transfected control mimics or mimics containing either the miR-124-3p wild-type (mimic-124) or the miR-124-3p seed mutant (mimic-124mut) in Huh7.5.1 cells, before infecting them with SINV-GFP (Figure 2A). Viral titer quantification demonstrated that the positive effect of miR-124-3p expression was lost when cells were transfected with the seed mutant mimic (Figure 2B). The same effect was observed on the viral capsid protein production (Figure 2C).

**Figure 2.**
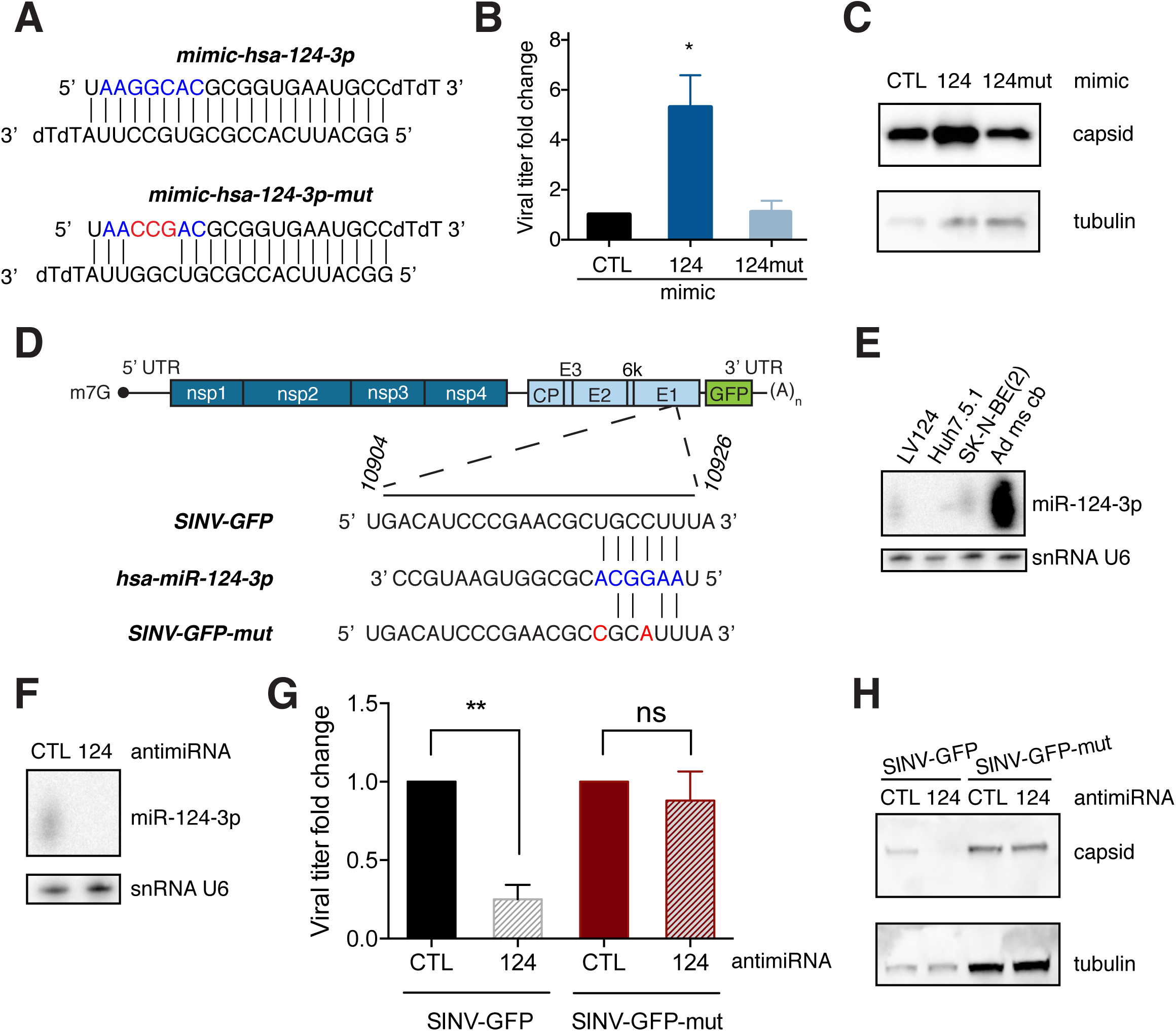
Importance of miR-124-3p seed in the positive regulation of SINV infection. (**A**) Diagram of mimic-124 and mimic-124-mut seed mutant. Seed sequence is indicated in blue and mutations in red. (**B**) Plaque assay quantification of SINV-GFP viral titers and (**C**) western blot analysis of viral capsid protein accumulation after transfection of mimic-124 and mimic-124-mut in Huh7.5.1 cells. Tubulin is used as a loading control (**D**) Schematic representation of SINV-GFP genomic structure with a diagram of miR-124-3p binding site within glycoprotein E1 coding region. Point mutations introduced in the mutant virus (SINV-GFP-mut) are indicated in red. (**E**) miR-124-3p expression in lentiviral transduced Huh7.5.1 cells stably expressing miR-124-3p (LV124), wild-type Huh7.5.1 cells, SK-N-BE(2) neuroblastoma cells and in adult mouse cerebellum (Ad ms cb). (**F**) Northern blot analysis of miR-124-3p expression in LV124 after antimiRNA transfection. snRNA U6 is used as a loading control. (**G**) Fold difference in viral titers produced by SINV-GFP or SINV-GFP-mut in LV124 after antimiRNA transfection. Error bars represent mean ± SEM of three independent experiments, ** p < 0.01, ns: non-significant, unpaired Student’s *t* test. (**H**) Western blot analysis of viral capsid protein accumulation in LV124 after antimiR transfection. Tubulin is used as a loading control.

As miRNA-mediated viral regulation can happen by direct binding of the miRNA to the viral RNA (33), we bioinformatically searched for predicted miR-124-3p binding sites on SINV genome. Using the prediction tool ViTa (34) we found three predicted binding sites at positions 2454-2475, 3828-3853 and 10904-10926 in the viral genome. We further studied the latter, localized within the virus ORF2 and more specifically in the glycoprotein E1 coding region (Figure 2D), which possesses a canonical 6-nt seed-match (nucleotides 2 to 7) for miR-124-3p. Using available SINV sequences on the NCBI Virus (35) database we sought to determine the conservation of this region among SINV strains. Sequence alignment of the entire genomic sequence of sixteen SINV strains isolated world-wide from various insects and vertebrates between 1969 and 2016 compared to the reference SINV genome (NC_001547) revealed that most of the strains (14 out of 16) share the same nucleotide composition at the binding site independently of the host, the year or the geographical areas in which they were isolated (Figure S4). Moreover, two of the most ancient and phylogenetically distant strains (MG182396 and KF981618) display silent point mutations in the miRNA binding site (Figure S4).

With the aim of disrupting the putative binding site without interfering with the viral coding sequence, we introduced two silent mutations at positions 10919 and 10922 (Figure 2D) in the viral genome. We first characterized the mutant virus (SINV-GFP-T10919C-C10922A, later on referred to as SINV-GFP-mut) by an infection kinetic study compared to the wildtype SINV-GFP in Huh7.5.1 cells. SINV-GFP-mut growth kinetics showed no significant differences in viral titers compared to the wild type SINV-GFP (Figure S5A). Furthermore, we assessed the release and spread of the mutant virus by verifying the plaque size phenotype and we observed no difference in plaque size between the two viruses (Figure S5B).

In order to measure the effect of miR-124-3p inhibition on the WT and mutant SINV, we established a Huh7.5.1 cell line stably expressing the human miR-124-1 gene (LV124 cells). miR-124-3p levels were verified in LV124 cells and compared to the expression in differentiated human neuroblastoma SK-N-BE(2) cells and adult mouse cerebellum, both used as positive controls. Huh7.5.1 cells were used as negative control. Northern blot analysis showed that miR-124 expression in the LV124 cells was similar to the expression level in differentiated neuroblastoma cells, known to express miR-124 (36) (Figure 2E).

Antisense oligonucleotides have been previously used as a strategy to sequester and block miRNAs (37). To test whether miR-124-3p inhibition reduced SINV infection, we transfected an antimiR-124 or a control antimiRNA into LV124 cells which resulted in a complete depletion of the mature miRNA as assessed by northern blot analysis (Figure 2F). We then evaluated the effect of miR-124 inhibition in this setup on SINV-GFP and SINV-GFP-mut infection. After antimiR-124 transfection, SINV-GFP viral titers as well as the capsid viral protein synthesis were significantly reduced (Figure 2G and H). In contrast, SINV-GFP-mut viral titers and protein levels remained unchanged upon miR-124-3p inhibition (Figure 2G and H). These results show that inhibition of miR-124-3p negatively regulates SINV infection and that mutations introduced in the binding site on the viral RNA abolish the miRNA-mediated regulation.

### miR-124-3p inhibition in differentiated neuroblastoma cells restricts SINV-GFP infection

Since miR-124-3p is a neuron-specific microRNA, we turned to a system where the miRNA was expressed endogenously to verify both the miRNA effect on SINV viral infection and the effect of virus infection on the miRNA expression. The human neuroblastoma SK-N-BE(2) cells can be differentiated into neuron-like cells by retinoic acid (RA) treatment (38). We first verified the neurite outgrowth which is a morphological hallmark of neuroblastoma cell differentiation *in vitro* (Figure 3A). To confirm the correct differentiation, we verified by RT-qPCR analysis that the expression of the neuron proliferation marker *MYCN* decreased after RA treatment (39) (Figure 3B). miR-124-3p endogenous expression increased at 6-days post differentiation compared to proliferating cells, but it was not affected by SINV infection (Figure 3C). We also assessed the effect of miR-124-3p inhibition on the production of SINV-GFP wild-type and mutant in these cells by transfecting miR-124-3p or control antimiR prior to differentiation and infection. Northern blot analysis of miR-124-3p expression confirmed the depletion of miR-124-3p in both mock and SINV-infected conditions (Figure 3D). While SINV-GFP wild-type viral titers as well as viral protein levels were reduced of about 50% in antimiR-124 transfected cells compared to control (Figure 3E and F), the mutation in the viral RNA (SINV-GFP-mut) abolishes the effect. These results show that inhibition of miR-124-3p negatively affects SINV-GFP viral infection in human neuronal differentiated cells and that this regulation involves the miR-124 binding site at position 10904-10926 in the viral genome.

**Figure 3.**
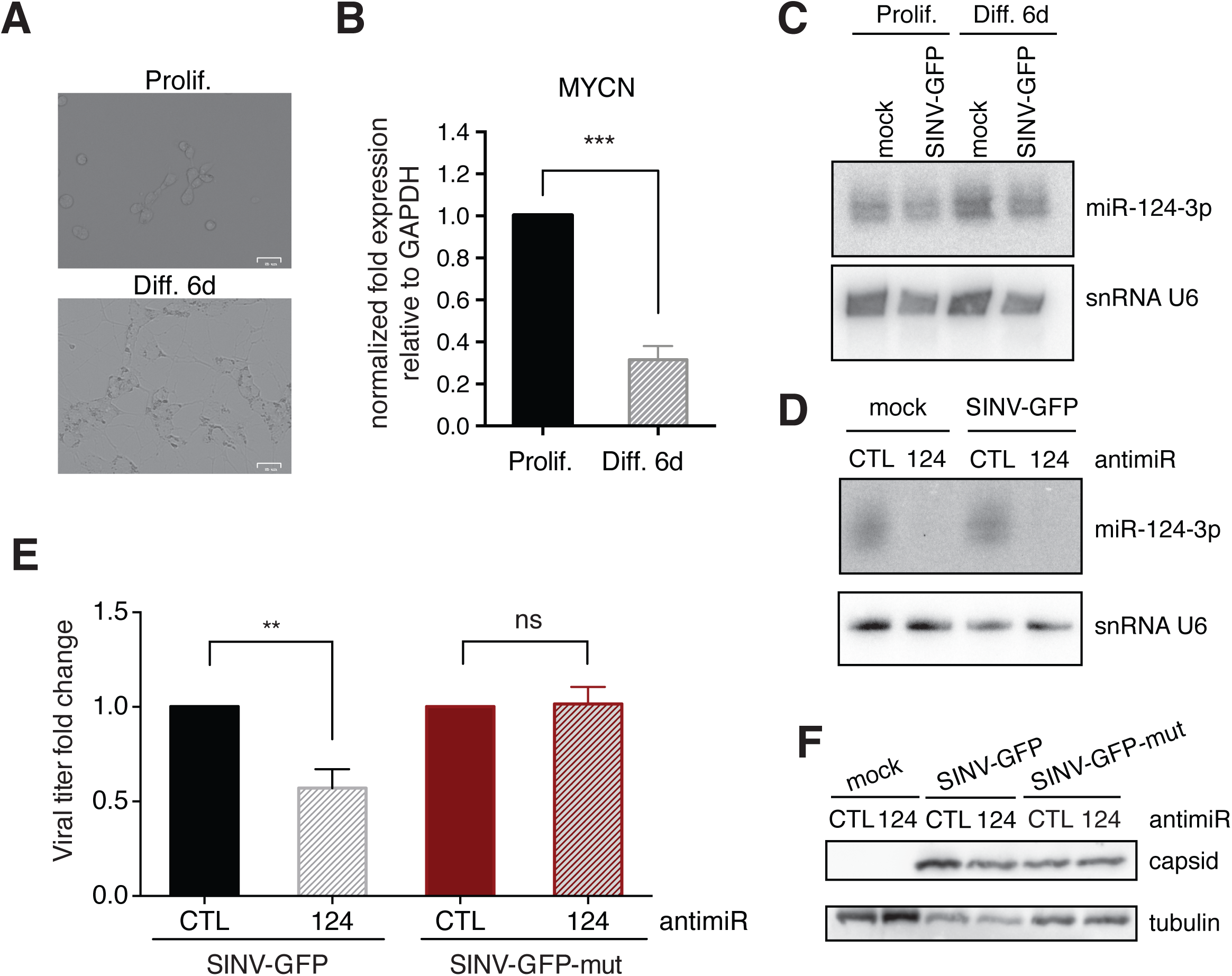
Sequestration of miR-124-3p in differentiated SK-N-BE(2) cells attenuates SINV-GFP infection. (**A**) Microscopy images of proliferating (upper panel) or 6 day-differentiated (lower panel) SK-N-BE(2) cells by 10 µM RA treatment. White bar corresponds to 25 µm (**B**) RT-qPCR analysis of relative *MYCN* expression in proliferating or differentiated cells. (**C**) Northern blot analysis of miR-124-3p expression levels in proliferating or 6 day-differentiated cells, mock or SINV-GFP infected. (**D**) Northern blot analysis of miR-124-3p expression levels upon antimiRNA transfection in differentiated cells at 6-day post RA treatment. (**E**) Plaque assay quantification of viral titer production of SINV-GFP or SINV-GFP-mut, shown as relative fold-change compared to control condition, and (**F**) capsid viral protein accumulation after antimiRNA transfection and 6-day post RA differentiation of SK-N-BE(2) cells. Tubulin is used as a loading control. Data are presented as mean ± SEM of three independent experiments. ** p < 0.01, *** p < 0.005, unpaired Student’s *t* test.

### Inhibition of miR-124-3p restricts CHIKV infection

In order to test whether miR-124-3p could positively regulate other positive, single-stranded RNA viruses, we tested two different alphavirus CHIKV strains (CHIKV La Réunion and Caribbean strains) and two strains of the flavivirus Zika virus (ZIKV African and French Polynesia strains). CHIKV La Réunion viral production was significantly increased following miR-124-3p overexpression. Though to a lesser extent, the viral production from the Caribbean strain also followed the tendency of increased titers following miR-124-3-p overexpression. In contrast, no significant effect could be observed on the two ZIKV strains tested (Figure 4A). To verify whether miR-124-3p inhibition could reduce CHIKV infection, we transfected an antimiR-124 or a control antimiRNA in LV124 cells and we infected them with a CHIKV La Réunion strain expressing GFP (Figure 4B). After antimiR-124 transfection, CHIKV-GFP viral titers were reduced of about 50% (Figure 4C). Moreover, antimiR-124 treatment significantly reduced CHIKV-GFP viral titers in differentiated neuroblastoma SK-N-BE(2) cells (Figure 4D). These results extend the proviral role of miR-124-3p to another alphavirus and confirm that miR-124-3p inhibition might be a promising strategy to block these viruses.

**Figure 4.**
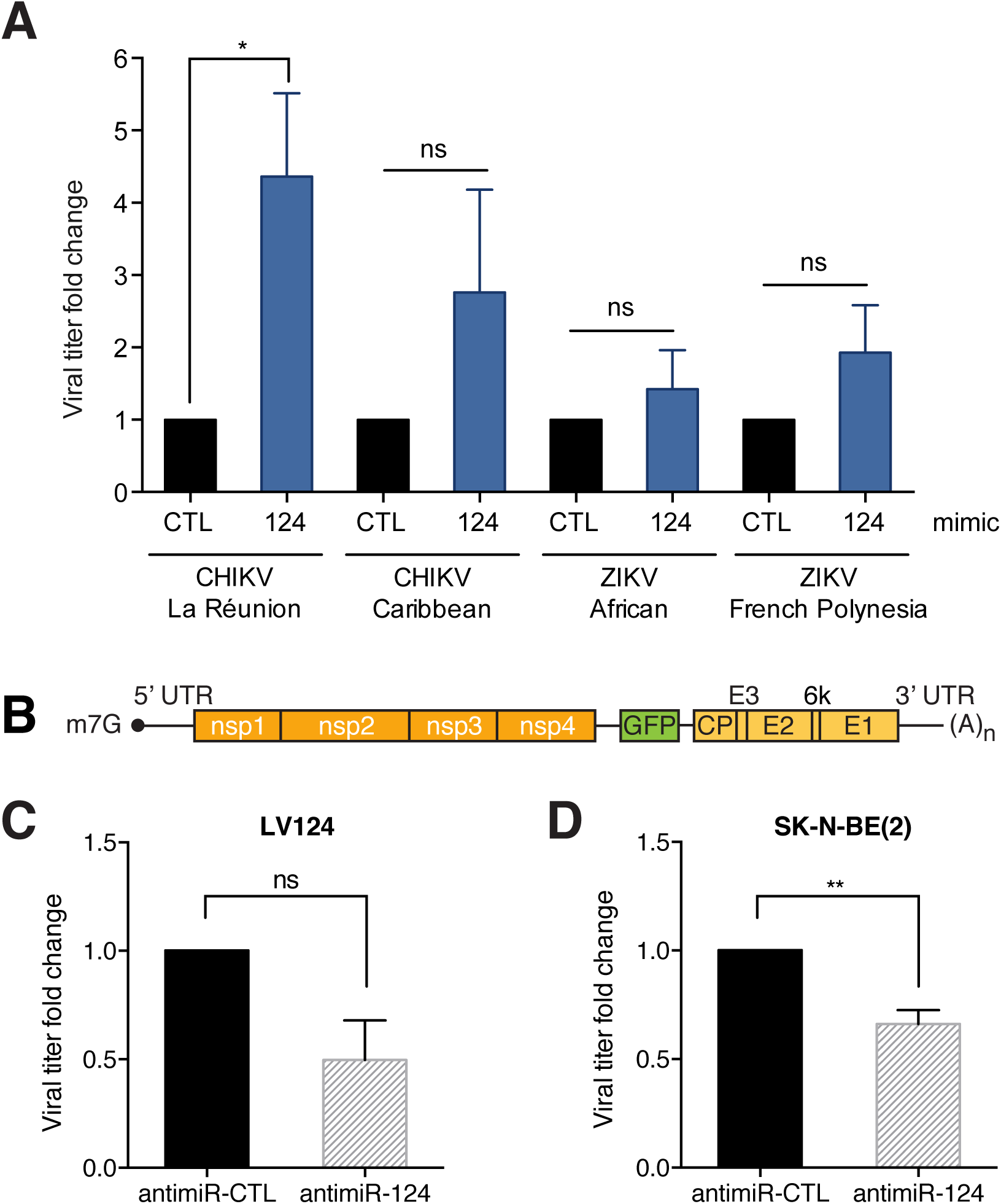
Sequestration of miR-124-3p in LV124 and differentiated SK-N-BE(2) cells attenuates CHIKV-GFP infection. (**A**) Plaque assay quantification of viral titers, shown as relative fold change, produced by CHIKV La Réunion strain, CHIKV Caribbean strain, ZIKV African strain and ZIKV French Polynesia strain after transfection of mimic-124-3p (blue bar) in Huh7.5.1 cells compared to control mimic (black bar). (**B**) Schematic representation of CHIKV-GFP genomic structure. (**C**) Plaque assay quantification of viral titer production of CHIKV-GFP in LV124 cells and (**D**) 6-days post RA differentiation in SK-N-BE(2) cells, shown as relative fold change of antimiR-124 compared to control condition. Data are presented as mean ± SEM of three independent experiments. * p < 0.05, ** p < 0.01, for (**A**) Two-way ANOVA with Bonferroni correction, for (**C**) and (**D**) unpaired Student’s *t* test.

## DISCUSSION

In addition to their role as fine-tuners of cellular functions, microRNAs are emerging as important regulators of host-pathogen interaction (33). They can regulate viral infection either directly, by targeting RNAs of viral origin, or indirectly, by targeting host RNAs, with a positive or negative outcome for the virus. Given the cell-specificity of certain miRNAs (40), the resulting interactions with viral RNAs can participate in determining the tissue-tropism of viral pathogens. For instance, stable expression of the liver-specific miR-122 increases HCV replication in non-hepatic cells (41). Another example is the inhibition of the Eastern equine encephalitis virus (EEEV) cycle by miR-142-3p which binds to the 3’ UTR of the virus in hematopoietic/myeloid cells (17).

SINV belongs to the alphavirus genus which includes viruses already emerged or with the potential to emerge as important human pathogens (8). Interestingly, as the alphavirus genome mimics the host messenger RNA and its replication takes place in the cytoplasm, the incoming viral RNA has the potential to directly interact with cellular miRNAs. In agreement, binding sites for cellular microRNAs were identified by AGO-CLIP within the alphavirus CHIKV, SINV and Venezuelan equine encephalitis virus (VEEV) genomes (15).

In this study, we took advantage of a functional genome-wide screen approach (42, 43) to identify pro- and anti-viral miRNA activity on SINV. The gain of function approach identified sixteen miRNAs whose overexpression had a positive effect on the virus. In contrast, our loss-of-function screen based on the usage of anti-microRNAs was not suitable to identify any phenotypic effect. This could reflect the fact that Huh7.5.1 cells lack miRNAs that could play a role in SINV infection and hence no miRNAs naturally expressed in those cells can have an antiviral effect on SINV. This in agreement with previously published results showing that the lack of miRNAs do not have a significant impact on several viruses (44). Among the hits, we uncovered hsa-miR-124-3p as a novel positive regulator of SINV and we demonstrated that its overexpression increases SINV viral production while mutations in its seed sequence abolish this effect. We identified and validated a binding site for hsa-miR-124-3p in the SINV genome within positions 10904 and 10926, which corresponds both to the 3’ non-coding portion of the genomic RNA and to the E1 glycoprotein coding region in the sub-genome.

As SINV may have a neuronal tropism (45), the binding of the cellular miRNA to the viral genome could provide an evolutionary advantage and could be considered as a novel example of co-evolution between the virus and its host. Indeed, we were able to show that miR-124 inhibitors reduced SINV infection in differentiated human neuroblastoma cells expressing miR-124-3p endogenously.

The importance of this miRNA/viral RNA interaction is supported by the evidence that SINV strains found in nature evolved to acquire and keep the miR-124 proper binding site while maintaining the E1 protein coding sequence.

Further work will be needed to determine the molecular mechanism involved in the positive regulation of the virus by miR-124-3p. Moreover, it will be of interest to study the contribution of the other two predicted miR-124 binding sites on SINV. At this stage, we cannot exclude that the SINV RNA serves as a miRNA sponge by sequestering miR-124 and derepressing pro-viral endogenous targets, as it is the case for Bovine Viral Diarrhea Virus and miR-17 (15) or HCV and miR-122 (46).

Finally, cellular miRNAs can also regulate viral infections by targeting cellular RNAs positively or negatively involved in the host response (33). Indeed, despite the absence of a predicted binding site on the viral genome, we also described a pro-viral effect of miR-124 on another medically relevant alphavirus. Inhibition of miR-124 expression reduces chikungunya virus (CHIKV) viral production in human cells independently of the direct binding to the viral RNA, supporting the idea that miR-124 may also play a role in the regulation of cellular targets against alphaviruses. This has already been observed for miR-124 regulation of measles virus (47), JEV (48) or HIV (49) infections. In addition, miR-124 expression is modulated by different viruses including ZIKV (50), EV71 (51), HCMV (52) or influenza H1N1 (53) suggesting a possible implication of the miRNA in the regulation of these infections as well. CHIKV and SINV are classified as arthritogenic alphaviruses (5) due to the natural symptoms of the infection, which include arthritis and bone pathology (54). This suggests that prior to reaching the CNS, productive infection of cell types other than neurons must take place. Interestingly, in addition to its involvement in the CNS, miR-124 has more recently emerged as a critical modulator of immunity and inflammation (55), by preventing microglia activation (56) or by regulating the adaptive immune response through STAT3 regulation (57). Furthermore, it has also been linked to bone pathology by playing a role in the regulation of osteoclast differentiation (58, 59). Thus, we cannot exclude a role of miR-124/SINV interaction in these cell types. It would be of interest to study miR-124 expression during alphavirus infection in synovial tissues and cartilage, the latter being more physiologically relevant cell types according to these viruses’ tropism. Indeed, this could give more insight into a possible advantage of miR-124/SINV interaction in establishing alphavirus-induced arthritis rather than encephalitis.

Developing therapeutic drugs to treat clinical diseases caused by previously neglected emerging viruses is therefore crucial for public health. Based on our work, we foresee the usage of miRNA inhibitors as a promising strategy in the fight against alphaviruses in the future. Finally, given the extensive usage of alphaviruses as a vehicle for vaccine and gene-therapy delivery (60), the identification of a positive regulation by miR-124 may have a strong impact on the development of more powerful biotechnologies based on SINV genome.

## Materials and methods

### Viral stocks, cell culture, and virus infection

Plasmids carrying a green fluorescent protein (GFP)-SINV genomic sequence or a green fluorescent protein (GFP)-CHIKV La Réunion genomic sequence (kindly provided by Dr. Carla Saleh, Institut Pasteur, Paris, France) were linearized with XhoI and NotI, respectively as in (61). They were used as a substrate for *in vitro* transcription using mMESSAGE mMACHINE capped RNA transcription kit (Ambion, Thermo Fisher Scientific Inc.) following the manufacturer’s instructions. GFP expression is driven by duplication of the sub-genomic promoter. SINV-GFP and CHIKV-GFP viral stocks were prepared in BHK21 baby hamster kidney cells, and titers were measured by plaque assay. The CHIKV strains used are La Reunion, 06-049 AM258994 and Caribbean (62). The ZIKV strains used are the African HD78788 and French Polynesia, PF-13.

The SINV-GFP-mut plasmid was generated by site directed mutagenesis using forward (5’ CGCATTTATCAGGACATCAGATGCACCACTGGTCTCA 3’) and reverse (5’ ATGTCCTGATAAATGCGGCGTTCGGGATGTCAATAGA 3’) primers and the In-Fusion HD Cloning Kit (Takara) following the manufacturer’s instructions.

Cells were infected with all viruses at a MOI of 10^−3^ and samples were harvested at 24 hours post-infection (hpi) unless specified otherwise.

Huh7.5.1 cells were maintained in Dulbecco’s Modified Eagle Medium (DMEM) 4.5 g/L glucose (Gibco, Thermo Fisher Scientific Inc.) supplemented with 10% fetal bovine serum (FBS) (Takara), 1% MEM Non-Essential Amino Acids Solution (NEAA 100X, Gibco, Thermo Fisher Scientific Inc.) and Gentamicin (50 µg/mL) (Gibco, Thermo Fisher Scientific Inc.) in a humidified atmosphere of 5% CO_2_ at 37°C. HEK293A (QBiogene) and Vero R (88020401, Sigma-Aldrich) cells were maintained in DMEM (Gibco, Thermo Fisher Scientific Inc.) supplemented with 10% FBS (Clontech) in a humidified atmosphere of 5% CO_2_ at 37°C.

SK-N-BE(2) cells (95011815, Sigma-Aldrich) were maintained in 1:1 medium composed of Ham’s F12 medium (Gibco, Thermo Fisher Scientific Inc.) supplemented with 15% FBS and Eagle’s Minimal Essential Medium (ENEM) supplemented with 1% NEAA (Gibco, Thermo Fisher Scientific Inc.). For miR-124-3p inhibition experiments in SK-N-BE(2) cells, due to the low transfection efficiency of these cells once differentiated, they were first transfected with 75 nM of antimiR specific to miR-124-3p or antimiR-CTL and 6 h later, differentiation was induced by 10 µM RA treatment (R2625, Sigma-Aldrich).

### Lentivirus production and generation of stable cell line

The human pre-miR-124a-1 gene in locus 8p23.1 flanked by about 200 nucleotides of its upstream and downstream genomic sequences was PCR amplified using the following primers: hsa-pri-miR-124 Fw 5’ GGGGACAAGTTTGTACAAAAAAGCAGGCTTCGAGCTGCGGCGGGGAGGATGC 3’; hsa-pri-miR-124 Rv 5’ GGGGACCACTTTGTACAAGAAAGCTGGGTCCCCCTGTCTGTCACAGGCTGC 3’. The obtained insert was first cloned by gateway BP reaction (Invitrogen, Thermo Fisher Scientific Inc.) in pDONOR-221 and then by gateway LR reaction (Invitrogen, Thermo Fisher Scientific Inc.) in the lentiviral destination vector pLenti6.2-3xFLAG-V5-ccdB (87072, Addgene). Lentiviruses were produced by co-transfection of pLenti6.2-3xFLAG-V5-hsa-pri-miR-124 with packaging vectors encoding lentiviral gag and pol proteins, pPAX and Vesicular Stomatitis Virus (VSV) envelope glycoprotein, pVSV-G. 24 h post transfection, the supernatant was collected and used as inoculum for transduction. Huh7.5.1 cells were transduced with lentiviruses in the presence of polybrene (SC-134220, Santa Cruz Biotechnology) 4 µg/mL for 6 h. Afterwards, the inoculum was removed, and cells were incubated in complete medium. Selection pressure was applied by supplementing the complete medium with 15 µg/mL of blasticidin (Invivogen). Surviving cells were maintained in culture in the presence of blasticidin as a polyclonal miR-124-3p expressing cell line (LV124).

### High-content miRNA-based phenotypic screening

For the miRNA screen, the miRIDIAN microRNA Mimic (catalog number CS-001030) and Inhibitor (catalog number IH-001030) Libraries (19.0, human microRNAs) were purchased from Dharmacon (Thermo Fisher Scientific Inc.). For the mimic screen, 20 nM of each Mimic microRNA were transfected into Huh7.5.1 cells (cultured in DMEM 4.5g/L glucose, 10% hyclone FCS, 50 µg/mL gentamycine) grown in Greiner µClear 96-well microplates using a high-throughput (HT) reverse chemical transfection with the INTERFERin HTS delivery reagent (Polyplus-transfection SA, Illkirch France). For the inhibitor screen, 75 nM of each inhibitorm microRNA were transfected into Huh7.5.1 cells as described above. The HT transfection protocol was optimized for reaching 90-95% transfection efficiency with minimal toxicity on a TECAN Freedom EVO liquid handling workstation. The screens were performed in technical triplicates. To limit biological variability, cell passage (n=3 after thawing), serum batch and transfection reagent batch were strictly determined. Internal controls such as positive (siRNA targeting SINV 3’ UTR at nt 12345 and nt 12437: 5’ AACUCGAUGUACUUCCGAGGAUU 3’, Integrated DNA Technologies) and negative siRNA controls (ON-TARGETplus Non-targeting siRNA #2, Horizon Discovery, Dharmacon), transfection efficiency control (“PLK1” siRNA that leads to cell death) were added to each microplate to determine parameters for inter-plate and day-to-day variability. Three days post-transfection, the cells were subjected to SINV-GFP viral infection for 24 hours before fixation and staining with Hoechst 3342 (labeling nucleus compartment “Nuclei”). High-throughput cell imaging was carried out with the INCELL1000 HCS epifluorescent microscope to collect an average of ∼ 6,000 cells analyzed per microwell (Hoechst 3342 and GFP channels).

### Analysis of the high-content siRNA screening data

Hoechst 3342 and GFP signals were extracted for all individual cells using the Multi Target Analysis module of the INCELL1000 platform. These parameters describe the GFP-non-infected and GFP+ SINV infected Huh7.5.1 cell populations for each miRNA treatment. The robust Strictly Standardized Mean Difference (SSMD* or SSMDr) for each well value was calculated as described below according to (63).

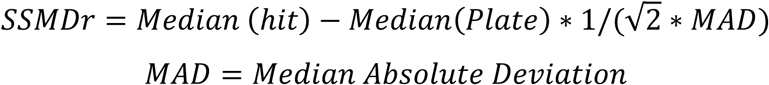

### Small RNA library preparation, sequencing and analysis

RNA was extracted from Huh7.5.1 cells and a small RNA library was prepared from 25 µg of total RNA as previously described in (61, 64, 65). Single-end sequencing was performed at the IGBMC Microarray and Sequencing platform, Illkirch, France on an Illumina Genome Analyzer IIx machine with a read length of 36 bp. Deep sequencing data analysis was performed as in (61) with slight modifications. Briefly, FASTX-Toolkit (http://hannonlab.cshl.eu/fastx_toolkit) was first applied to remove instances of the 3’ adaptor. Remaining reads between 18 and 32 nt in length were then mapped to the human genome (assembly version hg19 - UCSC repository), using Bowtie 1.0.0 (66). Up to 2 mismatches in total with no more than 1 mismatch in the first 15 nucleotides of each read were permitted. In addition, only alignments from the lowest mismatch-stratum were recorded and reads that could map to more than 50 loci were discarded. Finally, expressed human miRNAs (miRBase Release 20 (67)) were identified and quantified using BEDTools 2.16.2 (68) by comparing their genomic coordinates to those of the aligned reads. During the quantification process, multiple mapped reads were weighted by the number of mapping sites in other miRNAs and the final counts were normalized per million miRNA reads (RPM).

### Data accession number

The data discussed in this publication have been deposited in NCBI’s Gene Expression Omnibus (69) and are accessible through GEO Series accession number GSE136740.

### microRNA mimic and siRNA transfection

For reverse transfection in Huh7.5.1 or HEK293A cells, transfection complexes were prepared using mimics (miRIDIAN microRNA Mimics, Dharmacon) or siRNAs (Dharmacon or Integrated DNA Technologies) at a final concentration of 30 nM in INTERFERin^®^-HTS transfection reagent (Polyplus-transfection SA, Illkirch France), and transfection medium according to the manufacturer’s instructions. Transfection complexes were added to 1.5×10^4^ cells in each well of 48-well cell culture plates. Transfected cells were subsequently incubated for 72 hours before being infected with SINV-GFP at MOI 10^−3^ for 24 hours. For the experiment with the mutant mimic of miR-124-3p, mimics for *C*. *elegans* miR-67 (5’ UCACAACCUCCUAGAAAGAGdTdT 3’), miR-124-3p (5’ UAAGGCACGCGGUGAAUGCCdTdT 3’) and miR-124-3p mutated in the seed region (5’ UAACCGACGCGGUGAAUGCCdTdT 3’) were purchased at Integrated DNA Technologies.

### miRNA inhibition with 2′*O*-methylated oligonucleotides

For inhibition of miRNAs with antimiRs, 1.5×10^4^ LV124 cells were cultured in 48-well dishes and SK-N-BE(2) cells were cultured in 6-well plates at a confluency of 2×10^4^ cells/cm^2^. Cells were transfected with the 2′*O*-me antimiRs against the endogenously expressed miR-124-3p (5’ GGCAUUCACCGCGUGCCUUA 3’) or with the control sequence of *C*. *elegans* cel-miR-67 (5’ UCACAACCUCCUAGAAAGAGUAGA 3’), using INTERFERin^®^-HTS transfection reagent (Polyplus-transfection SA, Illkirch France) or Lipofectamine 2000 (Invitrogen, Thermo Fisher Scientific Inc.). AntimiRs were used at a final concentration of 75 nM and transfections were performed according to the manufacturer’s instructions. 48 h post-transfection cells were infected with SINV-GFP at a MOI of 10^−3^ for 24 h.

### Standard plaque assay

Vero R cells seeded either in 96- or 24-well plates were infected with infection supernatants prepared in cascade 10-fold dilutions for 1 h. Afterwards, the inoculum was removed and cells were cultured in 2.5% carboxymethyl cellulose for 72 h at 37°C in a humidified atmosphere of 5% CO_2_. Plaques were counted manually under the microscope. For plaque visualization, the medium was removed, cells were fixed with 4% formaldehyde for 20 min and stained with 1x crystal violet solution (2% crystal violet (Sigma-Aldrich), 20% ethanol, 4% formaldehyde).

### Western blotting

Proteins were extracted by collecting cell lysates in RIPA buffer (10 mM Tris/HCl pH 7.5, 150 mM NaCl, 0.5 mM EDTA, 0.1% SDS, 1% Triton X-100, 1% Sodium deoxycholate and protease inhibitor). Lysates were cleared by centrifugation at 13000 rpm for 30 min at 4°C to remove cell debris and the supernatant was retained for western blotting. Samples were loaded in a 10% acrylamide/bis-acrylamide gel and proteins were separated by migration at 100 V in 1x Tris-Glycine-SDS buffer. Proteins were transferred to a nitrocellulose membrane by wet transfer in 1x Tris-Glycine and 20% ethanol buffer. Viral proteins were detected using primary polyclonal antibodies against SINV CP (kind gift from Dr. Diane Griffin, Johns Hopkins University School of Medicine, Baltimore, MD) and a secondary antibody anti-rabbit-HRP (NA9340, GE Healthcare, Thermo Fisher Scientific Inc.). The signal was revealed by incubating the membrane for 10 min with SuperSignal West Femto Maximum Sensitivity Substrate (Pierce, Thermo Fisher Scientific Inc.). Tubulin was detected with a primary monoclonal antibody (T6557, Sigma-Aldrich) and a secondary antibody anti-mouse-HRP (NXA931, GE Healthcare Thermo Fisher Scientific Inc.).

### Northern blotting

Total RNA was extracted from cells with TRIzol Reagent (Invitrogen, Thermo Fisher Scientific Inc.) according to manufacturer’s instructions. 5-10 µg of total RNA were loaded on a 17.5% acrylamide-urea 4M gel and resolved by running in 1x Tris-Borate-EDTA for isolation of small RNAs. Nucleic acids were transferred to a nylon membrane by semi-dry transfer. Small RNAs were crosslinked to the membrane by chemical crosslink using N-(3-Dimethylaminopropyl)-N’-ethylcarbodiimide hydrochloride (EDC, Sigma-Aldrich). Membrane was pre-hybridized for 20 min with PerfectHyb(tm) Plus Hybridization Buffer (Sigma-Aldrich). DNA oligos directed against hsa-miR-124-3p (5’ GGCATTCACCGCGTGCCTTA 3’), hsa-miR-129-5p (5’ GCAAGCCCAGACCGCAAAAAG 3’), hsa-miR-665 (5’ AGGGGCCTCAGCCTCCTGGT 3’), hsa-miR-1244 (5’ AACCATCTCATACAAACCAACTACTT 3’), hsa-miR-30b-3p (5’ AGCTGAGTGTAGGATGTTTACA 3’) and snRNA U6 (5’ GCAGGGGCCATGCTAATCTTCTCTGTATCG 3’) were radiolabeled with 2.5 µCi of γ-ATP by PNK. After removal of unbound γ-ATP by MicroSpin G-25 column (GE Healthcare, Thermo Fisher Scientific Inc.) purification, the probe was incubated with the membrane in hybridization buffer overnight at 50°C. Membranes were washed twice with SSC 4x solution at 50°C and exposed on an image plate in a cassette. Imaging of the signal was obtained with Typhoon FLA 7000 laser scanner (GE Healthcare Life Sciences).

### RT-qPCR

DNase I treatment (Invitrogen, Thermo Fisher Scientific Inc.) was performed on 1 µg of RNA which was retrotranscribed using a random nonameric primer and the SuperScript IV Reverse Transcriptase (Invitrogen, Thermo Fisher Scientific Inc.) according to the manufacturer’s instructions. Quantitative PCR of *MYCN* expression was performed on 1:10 of cDNA using the SYBR Green PCR master mix (Thermo Fisher Scientific Inc.) with *MYCN* forward (5’ GAGCGATTCAGATGATGAAG 3’) and reverse (5’ TCGTTTGAGGATCAGCTC 3’) primers (39) or GAPDH forward (5’ CTTTGGTATCGTGGAAGGACT 3’) and reverse (5’ CCAGTGAGCTTCCCGTTCAG 3’) primers.

## ACKNOWLEDGMENTS

We thank our collaborators Carla Saleh who kindly provided us with SINV-GFP and CHIKV-GFP vectors, Diane Griffin for the SINV antibodies, Olivier Petitjean for producing the hsa-pri-miR-124-3p pDONR221 vector and Sophie Reibel-Foisset (Chronobiothron, UMS 3415) for the access to the BSL-3 facilities. The pLenti6.2-3xFLAG-V5-ccdB was a kind gift from Susan Lindquist (Addgene, #87072) (70). Sequencing was performed by the GenomEast platform, a member of the ‘France Génomique’ consortium (ANR-10-INBS-0009).

## FUNDINGS

This work was funded by the European Research Council (ERC-CoG-647455 RegulRNA) and was performed under the framework of the LABEX: ANR-10-LABX-0036_NETRNA, which benefits from a funding from the state managed by the French National Research Agency as part of the Investments for the future program. This work has also received funding from the People Programme (Marie Curie Actions) of the European Union’s Seventh Framework Program (FP7/2007-2013) under REA grant agreement n. PCOFUND-GA-2013-609102, through the PRESTIGE program coordinated by Campus France (to EG).

## AUTHOR CONTRIBUTIONS

SP and EG conceived the project. SP, EG and PL designed the work and analyzed the results. PL, EG, AW, BCM, PK, MM performed the experiments. EG and AW set up the high-throughput screen, AW performed the high-throughput screen and data acquisition, AK performed the bioinformatic analysis of the screen. BCWM generated the screen dot plot and table. PL and BCWM analyzed the microscopy images with CellProfiler software of selected candidates. PK generated the Huh7.5.1 small RNA libraries and BCWM performed the bioinformatic analysis. AF participated to the initial phase of the study for the candidate validation, PL and EG validated the mimic candidates. PL established the miR-124 stable cell line, generated and characterized the SINV-GFP mutant virus and produced CHIKV-GFP virus. PL and EG performed the miR inhibition assay with SINV-GFP and CHIKV-GFP in LV124 cells. BCM performed the ZIKV and CHIKV infections in mimic transfected Huh7.5.1 cells. EG, PL, BCWM and DBB identified miR-124 binding site. EG set up the SKNBE(2) differentiation experiments. PL and EG performed and analyzed SINV-GFP and CHIKV-GFP infections in differentiated SKNBE(2) cells, MM contributed to the time course experiments and western blot analysis. MV coordinated the work on ZIKV and CHIKV. LB coordinated the high-throughput screen and its bioinformatic analysis. PL and EG drafted the manuscript and designed the figures. PL, EG and SP wrote the manuscript with input from the other authors. SP and EG coordinated the work. SP assured funding. All authors revised the final manuscript.

