## Supplementary figures legends for "High-throughput fluorescence-based screen identifies the neuronal microRNA miR-124 as a positive regulator of alphavirus infection"

**Figure S1. Design and validation of siRNAs directed against SINV.**

(**A**) Schematic representation of SINV-GFP genomic structure with the corresponding position recognized by the siRNA SINV (siSINV) used as a functional control for the genome-wide screen. A repeated region within the 3’ UTR of SINV (starting at positions 12345 and 12437) is targeted. (**B**) Representative fluorescent microscopy images of Huh7.5.1 SINV-GFP infected after transfection with a control siRNA or with the siRNA directed against the virus (siSINV). (**C**) Percentage of GFP positive cells (left panel) and cell count to evaluate toxicity (right panel) after control siRNA (CTR1, CTR2, CTR4) or siSINV transfection. The average of at least 3 replicates is represented.

**Figure S2. Candidate microRNA expression in Huh7.5.1 cells and mouse adult tissues.**

(**A**) Top 30 microRNAs expressed in Huh7.5.1 cells. Ranking was based on the reads normalized per million microRNA reads (rpm) in the small RNA-seq experiment. (**B**) Northern blot analysis of miR-124-3p expression levels in adult and embryonic neuronal mouse tissues compared to heart and liver. (**C**) Northern blot analysis of miR-129-5p, miR-1244, miR-30b-3p and miR-665 in adult neuronal mouse tissues compared to heart and liver. Ethidium bromide (EtBr) staining was used as loading control.

**Figure S3. Validation of candidate mimics identified from the screen on SINV-GFP in HEK293A.** SINV-GFP fold-change in viral titers produced in HEK293A cells transfected with the top candidate mimics compared to the negative control (miR-137). Data correspond to three independent replicates plotted individually.

**Figure S4. Acquisition and conservation of miR-124-3p binding site in natural SINV isolates.** (**A**) Alignment of 16 SINV complete genomic sequences from natural isolates corresponding to nucleotide positions 10905 to 10925 in the reference genome (GeneBank accession n° NC_001547). Blue rectangle delimits predicted miR-124-3p binding site. Non-conserved nucleotides are highlighted in red (**B**) Phylogenetic tree generated from NCBI Virus sequence alignment of the entire genomic sequence of SINV strains in (A). In blue, the SINV reference sequence; in red, the isolates without miR-124-3p binding site conservation. For each strain, the GenBank accession number, host, geographical region and collection date are shown. Scale bar represents genetic distance.

**Figure S5. Characterization of the SINV-GFP-mut virus.** (**A**) Plaque assay quantification of viral titers produced by SINV-GFP and SINV-GFP-mut on Huh7.5.1 cells at 4, 8, 12 or 24 hpi. Error bars represent mean ± SD. ns, non-significant, Two-way ANOVA with Bonferroni correction. (**B**) Plaques for SINV-GFP and SINV-GFP-mut visualized by crystal violet staining on infected Vero cells to assess plaque-size phenotype.
