## Supplementary figures and images for "High-throughput fluorescence-based screen identifies the neuronal microRNA miR-124 as a positive regulator of alphavirus infection"

### Supplementary Figure S1

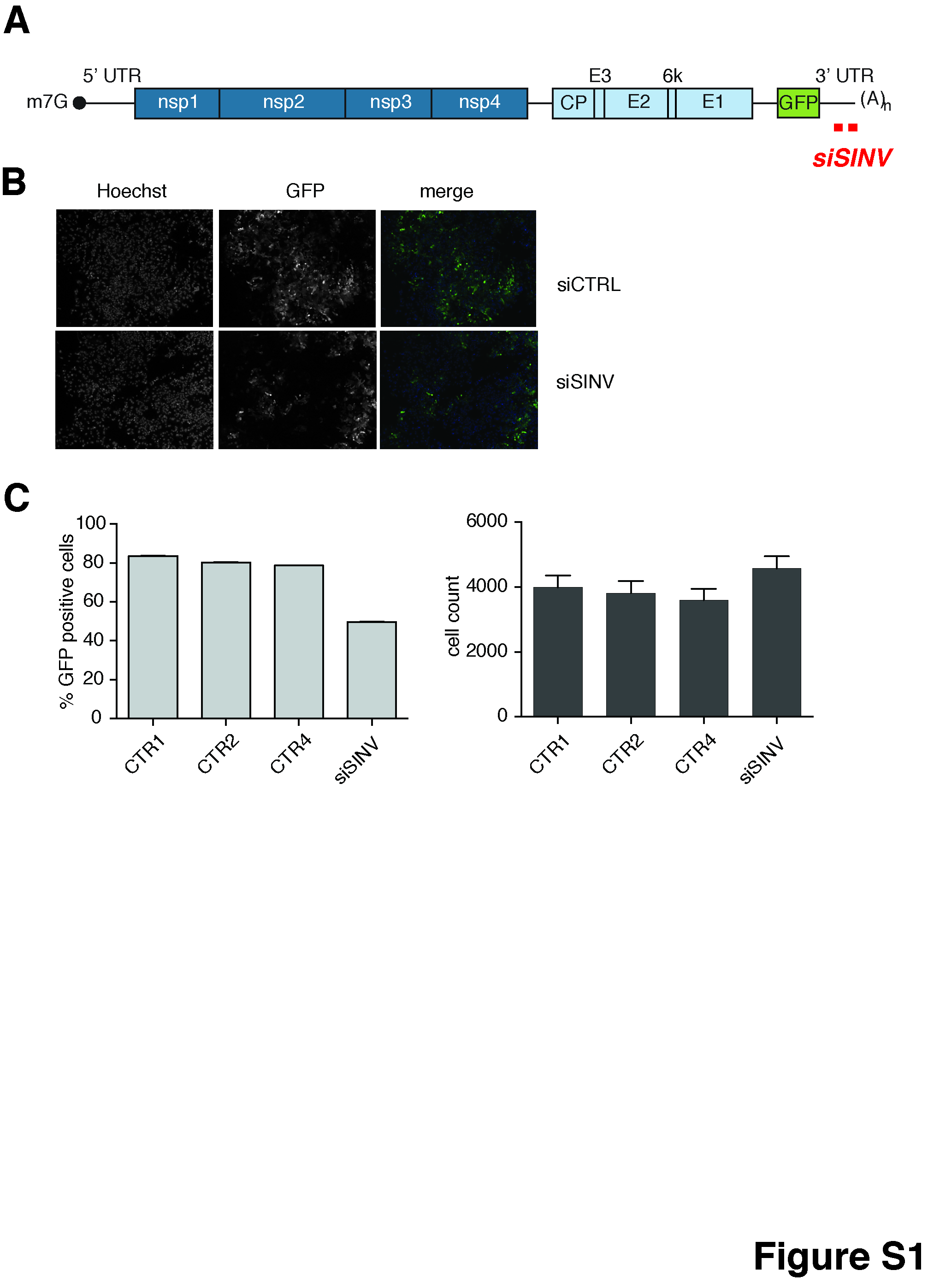

### Supplementary Figure S2

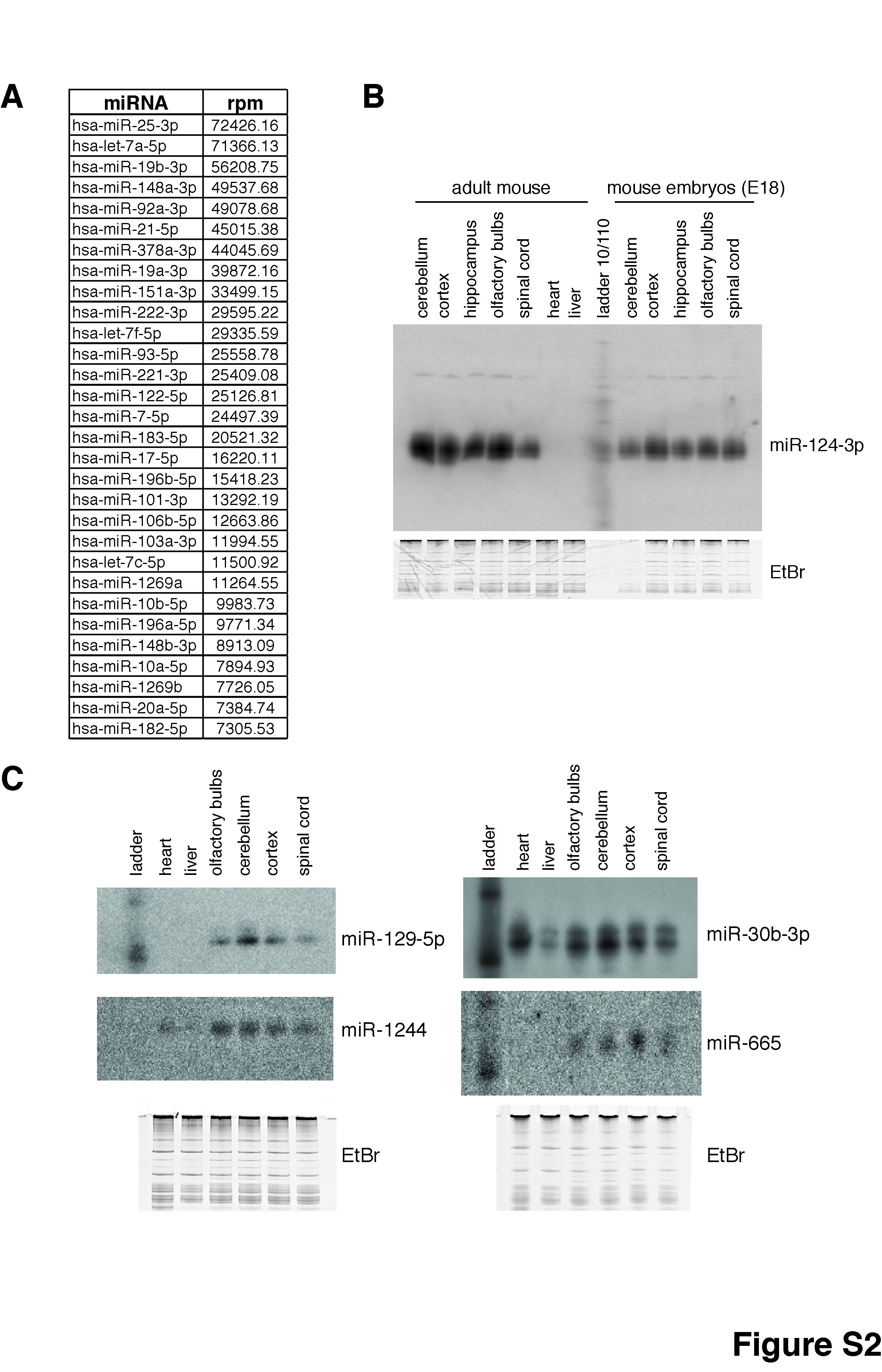

### Supplementary Figure S3

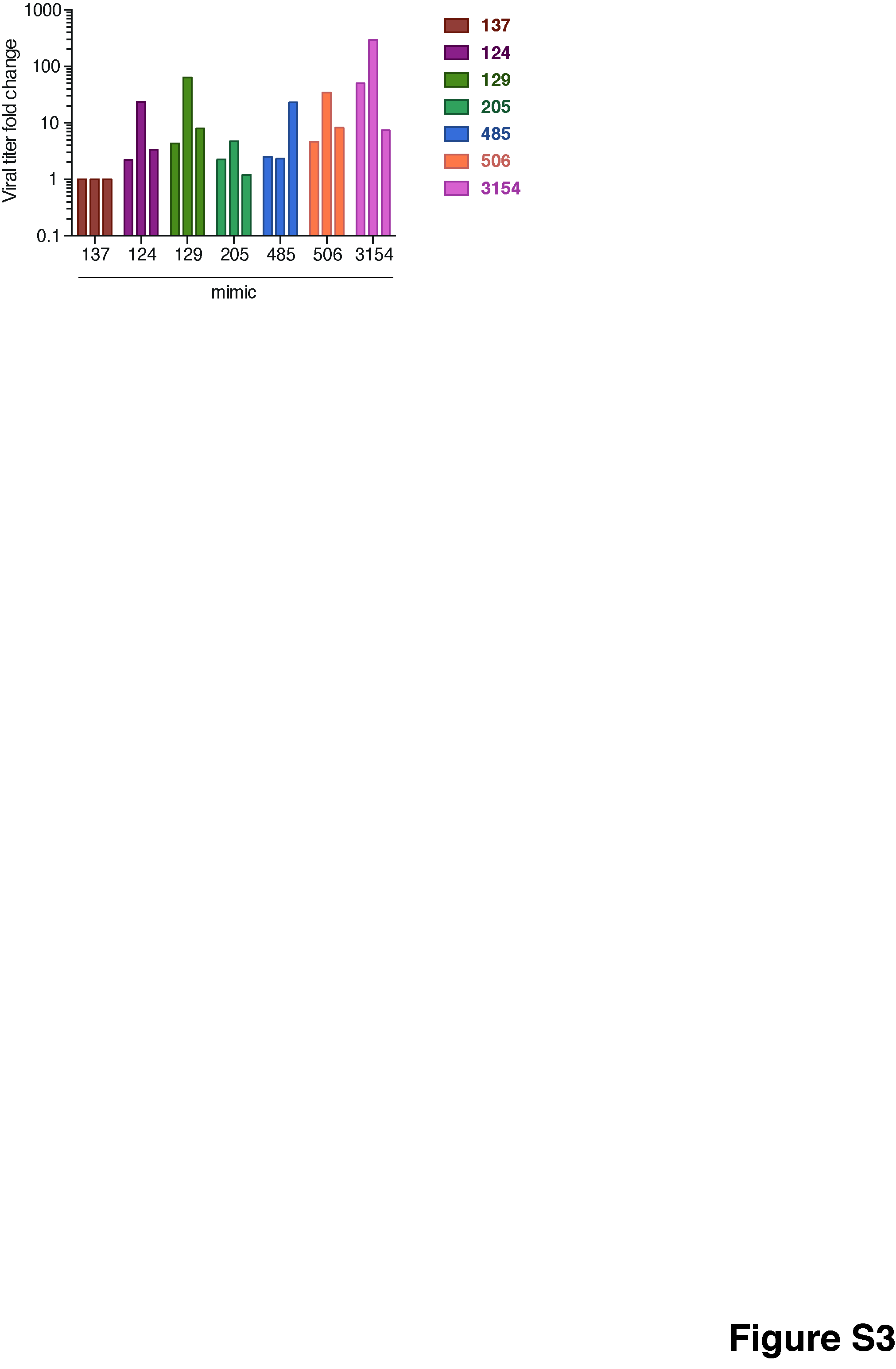

### Supplementary Figure S4

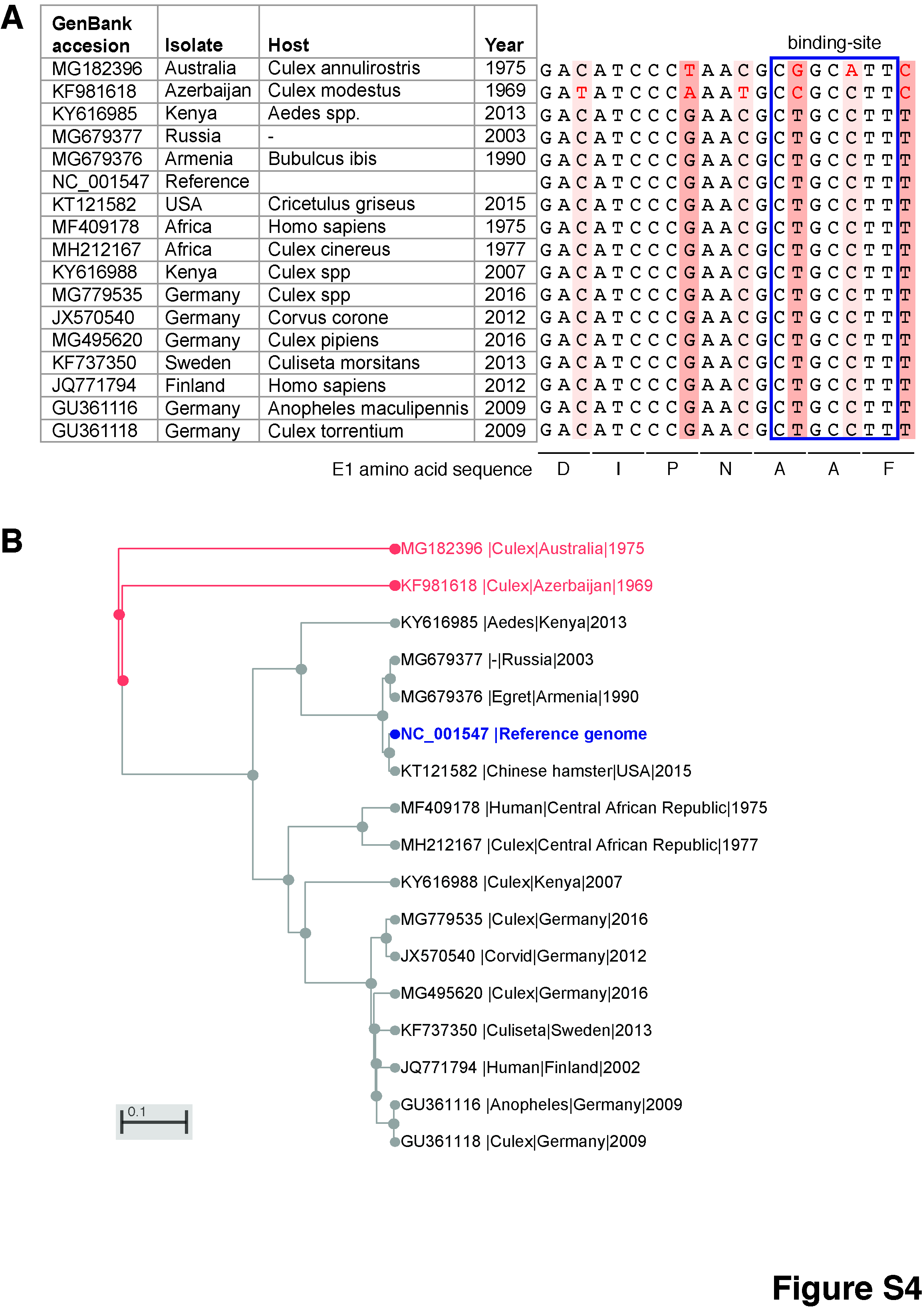

### Supplementary Figure S5

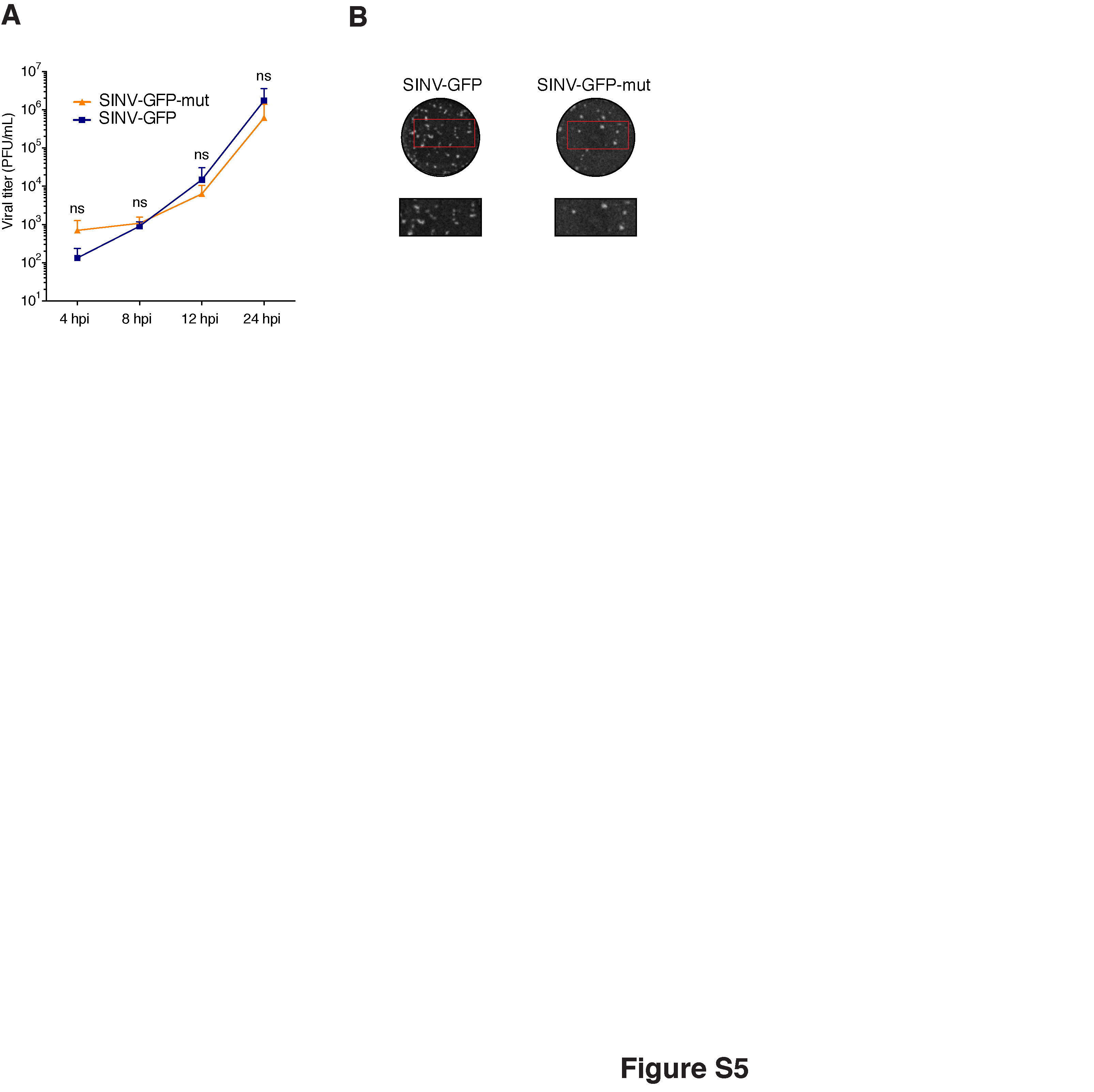
